# Integrative Transcriptomic Profiling of NK Cells and Monocytes: Advancing Diagnostic and Therapeutic Strategies for COVID-19

**DOI:** 10.1101/2024.10.21.619361

**Authors:** Salma Loukman, Reda Ben Mrid, Najat Bouchmaa, Hicham Hboub, Rachid El Fatimy, Rachid Benhida

## Abstract

In this study, we use integrated transcriptomic datasets from the GEO repository with the purpose of investigating immune dysregulation in COVID-19. Thus, in this context, we decided to be focused on NK cells and CD14+ monocytes gene expression, considering datasets GSE165461 and GSE198256, respectively. Other datasets with PBMCs, lung, olfactory, and sensory epithelium and lymph were used to provide robust validation for our results. This approach gave an integrated view of the immune responses in COVID-19, pointing out a set of potential biomarkers and therapeutic targets with special regard to standards of physiological conditions. IFI27, MKI67, CENPF, MBP, HBA2, TMEM158, THBD, HBA1, LHFPL2, SLA, and AC104564.3 were identified as key genes from our analysis that have critical biological processes related to inflammation, immune regulation, oxidative stress, and metabolic processes. Consequently, such processes are important in understanding the heterogeneous clinical manifestations of COVID-19—from acute to long-term effects now known as ‘long COVID’. Subsequent validation with additional datasets consolidated these genes as robust biomarkers with an important role in the diagnosis of COVID-19 and the prediction of its severity. Moreover, their enrichment in key pathophysiological pathways presented them as potential targets for therapeutic intervention.The results provide insight into the molecular dynamics of COVID-19 caused by cells such as NK cells and other monocytes. Thus, this study constitutes a solid basis for targeted diagnostic and therapeutic development and makes relevant contributions to ongoing research efforts toward better management and mitigation of the pandemic.

## 1 Introduction

COVID-19 is a viral infection caused by the SARS-CoV-2 virus, usually transmitted via respiratory droplets. The symptoms vary from mild to severe, and in serious conditions, it results in hospitalization, sometimes leading to death [1]. Despite the extensive research performed so far, diagnosis and management of COVID-19 remains tricky because there is no specific biomarker and the underlying pathophysiology is poorly understood. The transcriptomic analysis has become the tool of choice in identifying not only possible therapeutic targets but also biomarkers of various abnormalities, including COVID-19. [2]

Indeed, transcriptomic analysis has provided valuable information on the host immune response and dysregulated pathways and possible biomarkers in the context of COVID-19. Such comparison of transcriptomic studies between COVID-19 patients and healthy individuals or patients with other respiratory diseases has established different patterns of gene expression associated with SARS-CoV-2 infection. Several data published illustrated the dysregulation of immune-related genes, inflammatory responses, and coagulation pathways in COVID-19, which thus could play a major role in targeted treatments that could be developed based on this knowledge [3], [4], [5].

For example, transcriptomic studies have highlighted critical host factors in SARS-CoV-2 infection, including the ACE2 receptor and the TMPRSS2 protease, which can be targeted for therapeutic intervention [6]. Moreover, by using transcriptomic analysis, the assessment of treatment response analysis is greatly enhanced through the observation of changes in gene expression patterns after therapeutic interventions [7]. However, due to the lack of reliable biomarkers, COVID-19 remains poorly managed. Identifying specific and strong biomarkers of COVID-19 has been quite a challenge; this really requires an in-depth interaction between the virus and the host’s immunity, coupled with variations in immune and genetic makeups.

Recent advances in transcriptomics have highlighted the key role of selective immune cells, notably NK cells and CD14+ monocytes. These cells have emerged as vital players in the innate immune response, exhibiting specific expression profiles in COVID-19 patients that may mirror disease severity and clinical outcomes. This huge potential of cellular markers in the clinical set-up remains underutilized due to the complex nature of their modulation and interaction with viruses. [8], [9], [10], [11]

In this regard, we employed several extensive transcriptomic datasets from GEO for an in-depth understanding of immune dysregulation in COVID-19. More importantly, we looked into two unique datasets, GSE165461 and GSE198256, that provided specific insights into the gene expressions of NK cells and CD14+ monocytes in COVID-19 patients. GSE19825 allowed the investigation of longitudinal immune responses both in infection and recovery, while GSE165461 captured immune system behavior at different disease states. By identifying genes differentially expressed in these critical immune cells, we tried to uncover the mechanisms behind severe immune responses in COVID-19.

We have also added more datasets to reinforce the strength of our results for the GEO analysis. The additional datasets included various tissue types, including PBMCs, lung, OE, and lymph. Such a validation of results across wider biological contexts by integration of a broader dataset had allowed us to ensure that the analysis of gene expression changes was comprehensive. Single-cell studies, cell lines, and those involving pharmacological treatments were carefully avoided in order to maintain a focus on standard physiological responses. What’s more, this would definitely reinforce the reliability of our observations, since the holistic view on immune responses in COVID-19 provides a lot of potential biomarkers and therapeutic targets.

Our findings underline that a cluster of genes, including *SLA, CD163, MKI67, MAFG, ADAM19, IFI27, RNASE2, HBA1, HBA2, CENPF, TMEM158, THBD, S100A9, S100A8, and MBP*, show significant differential expression with key biological processes involved in inflammation, immune system modulation, oxidative stress, and metabolic activity. These genes, beyond their potential role as markers for early detection in COVID-19 infection, also have predictive value in the severity determination of the disease. *MBP, IFI27, MKI67, CD163, MAFG, and ADAM19* showed the highest expression pattern, with *SLA*, indicating progression toward acute respiratory complications and, therefore, might form the basis in planning individual treatment specifically.

More importantly, the genes shown in this study extend beyond the acute phase of the disease, where genes are still in altered expression levels well into the recovery phase. Surprisingly, *AC104564*.*3, HBA2, TMEM158, THBD, HBA1, ANKRD33B, LHFPL2, and SLA* were persistently overexpressed even at three months after recovery, thus being candidate genes that could be further studied as biomarkers for long COVID.

Value obtained from advanced transcriptomic technologies coupled with thorough data analysis provides an insight into the molecular signature of COVID-19 that was not available hitherto. Further, this work enhances the current scientific understanding and supports the development of targeted diagnostic tools and therapeutic strategies. Our findings may well influence how COVID-19 is currently being diagnosed, managed, and treated, bringing personalized medicine to the forefront in this battle against this health challenge.

## 2 Background & Summary

To detect SARS-CoV-2 variations big efforts have been deployed using genomic sequencing. Moreover, to examine changes in gene expression in COVID-19 patients, RNA sequencing (RNA-seq) assays has been used. Despite the wealth of OMICs data collected from COVID-19 patients over the past three years, the underlying mechanisms of the disease remain poorly understood[12]. Moreover, many individuals continue to experience long-term COVID-19 effects, highlighting the urgent need for a more systematic approach to data analysis, particularly RNA-seq data mining. Through the examination of shared differentially expressed genes, we successfully pinpointed novel potential biomarkers for diagnosing COVID-19 and identified potential therapeutic targets. The design of the study is presented in Figure-1

**Figure-1.**
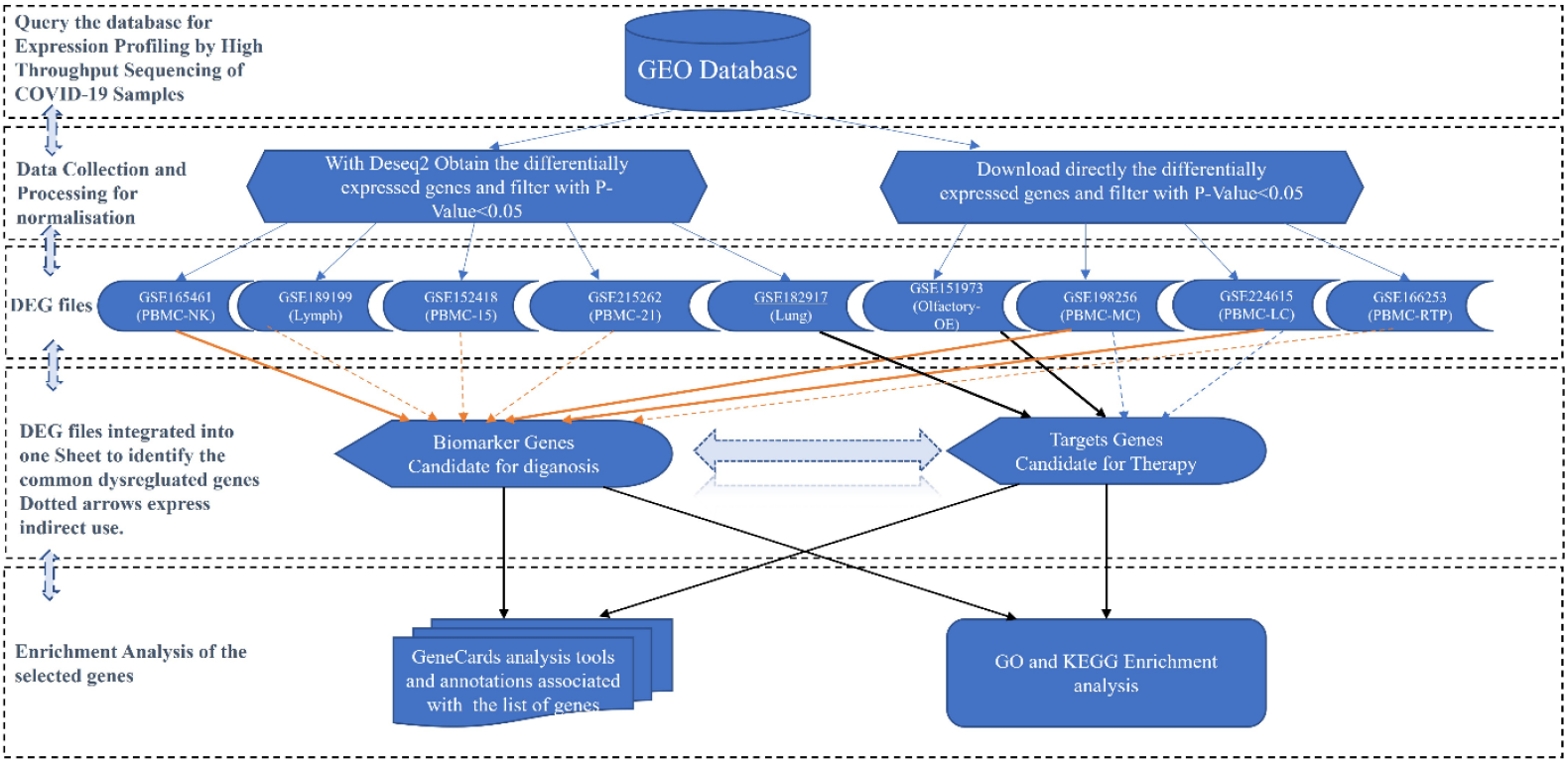
The workflow description: The workflow description of the different steps of the study, from the data collection to the analysis and information production. Dotted arrows indicate indirect use to support analysis.

## 3 Results and Discussion

### Identification of new biomarkers for the determination of the severity of COVID-19

The primary aim of this study was to identify molecular markers for the diagnosis of COVID-19. We hypothesized that transcriptomic data from blood samples could reveal biomarkers effective for the rapid diagnosis of the disease and its severity. We selected the datasets GSE165461 and GSE198256 for their potential insights into genes associated with SARS-CoV-2 infection.

In our analysis, we utilized GSE165461, which provides data on expression patterns of Natural Killer (NK) cells from both healthy donors and COVID-19 patients, across varying degrees of disease severity. Additionally, GSE198256 was employed to examine CD14+ monocyte expression in healthy donors, acute COVID-19 cases, and recovered patients. These datasets were chosen to give a comprehensive view of the immune response mechanisms disrupted by COVID-19.

The use of both datasets, GSE165461 and GSE198256, as starting points is justified by the comprehensive insights they provide into the immune responses crucial for combating COVID-19. In fact, NK cells are integral to the innate immune defense, acting swiftly to destroy virus-infected cells and regulate immune responses through their cytokine production. Their dysfunction or depletion, particularly observed in severe COVID-19 cases, underscores their importance in disease progression and potential as therapeutic targets [22]. Similarly, CD14+ monocytes play a pivotal role in the inflammatory response to infection. Their ability to produce significant quantities of pro-inflammatory cytokines, while essential for fighting the virus, can also lead to the cytokine storm associated with severe disease outcomes. The heightened activity of these monocytes in severe cases highlights their utility as biomarkers for assessing disease severity [8], [9].

These datasets, GSE165461 and GSE198256, offer valuable RNA sequencing data that allow for the detailed analysis of these critical immune cells under various clinical conditions. GSE165461 provides a longitudinal perspective, capturing the dynamics of the immune response across different stages of COVID-19, which is crucial for understanding how these cellular mechanisms evolve during the disease. GSE198256, with its focus on specific immune cells like NK cells and CD14+ monocytes, enables a deeper examination of the molecular pathways that these cells modulate during infection. This targeted analysis aids in identifying key genetic expressions that can serve as reliable biomarkers for diagnosing and predicting the progression of COVID-19, thereby supporting the development of targeted therapies.

Both different datasets were employed to determine the differentially expressed genes (DEGs) using DESEQ2. The results obtained were filtered using a p-value <0.05. Venn diagram was used to obtain common genes between the both different datasets in different prism of view, and the results are presented in Figure-2.

**Figure-2.**
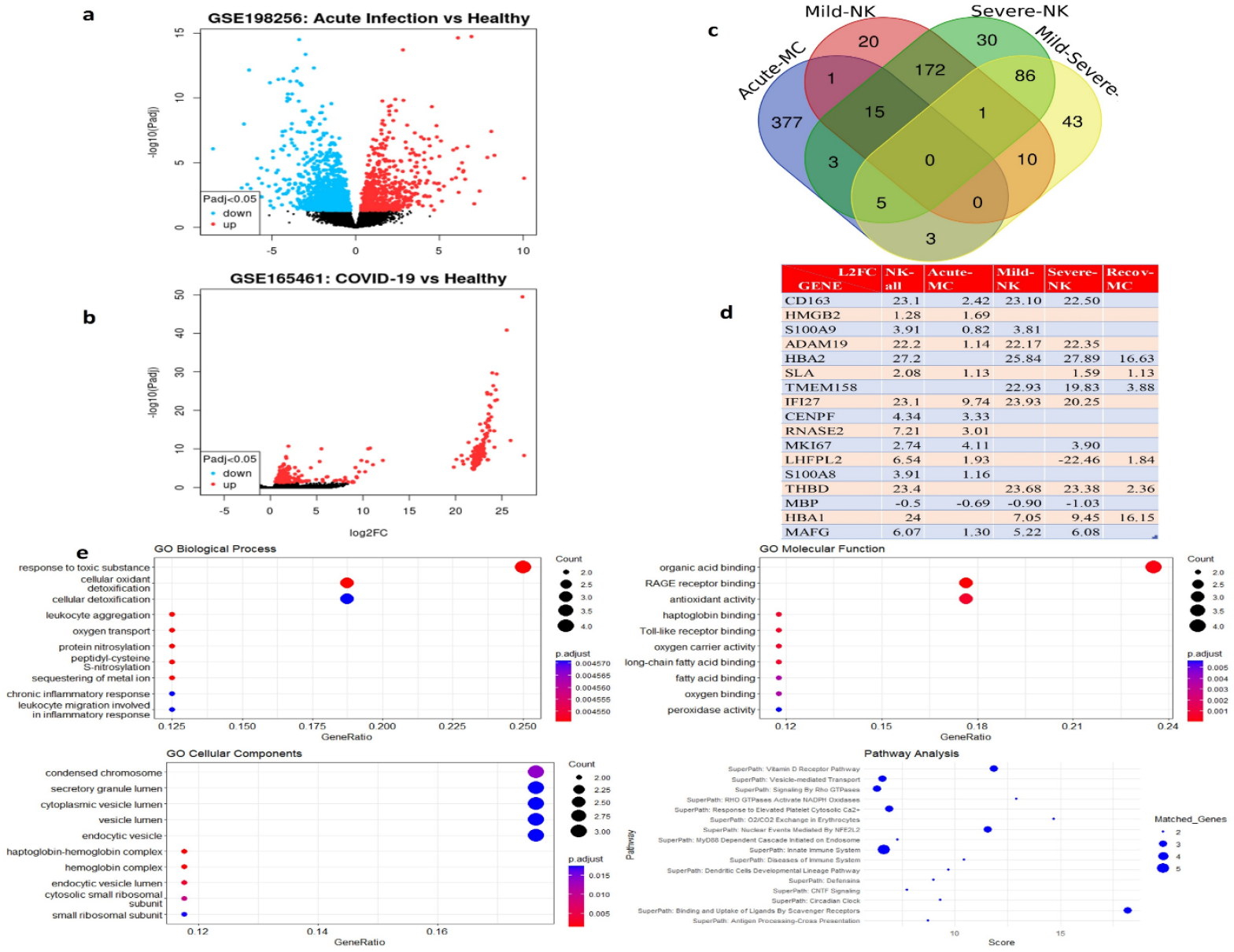
Transcriptomic profiling of COVID-19 diagnosis: a,b) Volcano plot of GSE165461 dataset (NK) and GSE198256 dataset (Acute-MC); c, Venn diagram of the differentially expressed genes between GSE165461 and GSE198256 datasets; d)a list of the common dysregulated genes in the two datasets GSE165461 and GSE198256; e) enrichment analysis of common dysregulated genes.

The results obtained shows that 17 genes are commonly dysregulated between the studied datasets, 16 of them are up-regulated including *SLA* (Src-like adaptor), *CD163* (CD163 molecule), *MKI67* (marker of proliferation Ki-67), *MAFG* (MAF bZIP transcription factor G), *ADAM19* (ADAM metallopeptidase domain 19), *IFI27* (interferon alpha-inducible protein 27), *RNASE2* (ribonuclease A family member 2), *HBA1* (hemoglobin subunit alpha 1), *HBA2* (hemoglobin subunit alpha 2), *CENPF* (centromere protein F), *TMEM158* (transmembrane protein 158), *THBD* (thrombomodulin), *S100A9* (S100 calcium binding protein A9), *HMGB2* (High Mobility Group Box 2), *LHFPL2* (LHFPL Tetraspan Subfamily Member 2) *and S100A8* (S100 calcium binding protein A8). Nevertheless, only *MBP* (Myelin Basic Protein) was commonly downregulated in these two datasets.

The 16 genes that are up-regulated have been identified as differentially expressed genes in various conditions and diseases, such as COVID-19. Upregulation of these genes has been associated with several functions and pathways, such as immune response, cell proliferation, and inflammation. Indeed, *MKI67, CENPF, RNASE2, HBA1, HBA2, TMEM158, THBD, MAFG, ADAM19, LHFPL2, and HMGB2* have been identified to be upregulated in other diseases such as cancer and autoimmune disorders[23], [24], [25], [26], [27], [28], [29], [30][31] Regarding SLA, it has been identified as a susceptibility gene for African swine fever virus (ASFV) infection in pigs and is involved in regulating immune responses [32]. While the specific roles of these genes and their impacts in COVID-19 have not been fully elucidated, upregulation of the genes *IFI27, CD163, S100A8, and S100A9* has been associated with COVID-19 pathogenesis[32], [33], [34]

Further research into the functions and effects of the newly discovered COVID-19 genes is therefore necessary, as their overexpression can have a variety of implications on the immune response, inflammation, tissue damage, and viral replication. Understanding their roles in the pathogenesis of COVID-19 could therefore offer new insights into the mechanisms underlying the disease and lead to the development of new therapeutic approaches, notwithstanding the interest in using them as biomarkers for identifying SARS-CoV-2 infection and severity.

A common down-regulated gene between the two dataset is the *MBP* gene which encodes for a specific protein within the myelin sheath of Schwann cells and oligodendrocytes[35]. Nevertheless, *MBP*-related transcripts were reported to be also expressed in the the immune system and bone marrow [36]. Myelin is a fatty substance that insulates and protects nerve fibers and allows for the efficient transmission of electrical impulses. In addition to its structural role, *MBP* is also thought to be responsible for myelination and maintaining the integrity of myelin during aging and disease. When the *MBP* gene is down-expressed, it can lead to a range of neurological disorders, particularly those affecting myelin function within the central nervous system (CNS).

No precise role has been attributed to *MBP* dysregulation in COVID-19, however, we suggest its implication in neurological symptoms and cognitive impairments that are observed in some COVID-19 patients, especially those with severe disease. More research is required to fully understand the mechanisms by which MBP dysregulation may contribute to the pathogenesis of COVID-19.

### Biological functions and signalling pathways of the selected genes

As earlier mentioned, the identified genes such as *SLA, CD163, MKI67, MAFG, ADAM19, IFI27, RNASE2, HBA1, HBA2, CENPF, TMEM158, THBD, S00A9, LHFPL2, HMGB2 and S100A8* have been found to be related to immune response, inflammation, and cell migration. We also used GeneCards online platform (Fishilevich et al., 2017) (https://ga.genecards.org/shared-query/d98759c6-236f-48a0-beb5-69ac7c025c42) to analyze in more details the functional relevance of these genes in relation to biological processes and pathways, molecular functions, and cellular components. The top biological processes associated with these genes include neutrophil aggregation, chemotaxis, sequestering of zinc ions, leukocyte aggregation, cellular oxidant detoxification, oxygen transport, protein nitrosylation, peptidyl-cysteine S-nitrosylation, sequestering of metal ions and chronic inflammatory response.

Neutrophil aggregation is the process of neutrophils binding together to form a cluster, which is a critical step in the innate immune response to invading pathogens [37]. Chemotaxis is the movement of cells towards a chemical gradient, such as a chemokine, which is important for leukocyte migration and immune cell recruitment to the site of infection[38]. Sequestering of zinc ions is a process that helps to limit the availability of zinc, which is an essential nutrient for invading pathogens [39].

The top relevant molecular functions associated with these genes are: organic acid binding, haptoglobin binding, Toll-like receptor binding, antioxidant activity, oxygen carrier activity, long-chain fatty acid binding, fatty acid binding, oxygen binding, and Microtubule binding. RAGE (Receptor for Advanced Glycation End-products) receptor binding was found to be among the most relevant molecular functions. RAGE is a cell surface receptor that is involved in many processes, including inflammation, oxidative stress, and immune cell activation [40]. Microtubule binding is important for many cellular functions, like intracellular transport, cell division and migration[41].

The top cellular components associated with these genes comprise the condensed chromosome, haptoglobin-hemoglobin complex, hemoglobin complex, endocytic vesicle lumen, cytosolic small ribosomal subunit, small ribosomal subunit and secretory granule lumen. The condensed chromosome is a tightly packed form of chromatin that is visible during cell division[42]. The haptoglobin-hemoglobin complex forms when haptoglobin binds to free hemoglobin, playing a vital role in preventing hemoglobin-induced oxidative damage and facilitating its clearance from the bloodstream [43]. Secretory granules are specialized vesicles that store and secrete proteins, such as cytokines and enzymes, which are implicated in multiple cellular processes that include inflammation and immune response[44].

The pathways identified in this analysis cover essential biological processes such as innate immunity, scavenger receptor-mediated ligand binding, vitamin D receptor signaling, and cellular stress responses mediated by NFE2L2. Additionally, pathways regulating responses to elevated platelet cytosolic calcium, vesicle-mediated transport, and Rho GTPase signaling were highlighted. These pathways play critical roles in cellular homeostasis, host defense, and cellular communication [45],[46].

In summary, the 17 genes identified in this analysis are related to several biological processes, molecular functions, and cellular components with a significant emphasis on immune system regulation, oxidative stress response, metabolic processes, inflammation, and cell migration. The analysis suggests that these genes could interfere in the regulation of these processes.

Considering the findings of the analysis that suggest a potential role for the identified genes in various biological processes, it is essential to investigate their expression patterns in specific cell types and disease conditions. In this regard, the expression of these genes in two datasets GSE165461 dataset (NK) and GSE198256 dataset (MC) could provide valuable insights into their involvement in the different conditions associated with COVID-19. In addition, the expression patterns of these genes in NK and MC cells could shed light on their potential as biomarkers for COVID-19 treatment.

### Expression patterns of the seventeen genes in NK cells and MC cells of COVID-19 patients

The study examined the expression distribution of the seventeen identified genes under different conditions across two datasets, GSE165461 dataset (NK) and GSE198256 dataset (MC). Boxplots in Figure-3 display the expression levels of these genes in various groups within each dataset. The GSE198256 dataset (MC) consisted of three groups: healthy controls, acute patients and recovered individuals after three months. On the contrary, the GSE165461 dataset (NK) comprised three groups: healthy controls, patients with mild symptoms, and patients with severe symptoms. Among the genes analyzed, eight emerged as potential predictors of disease severity. These findings suggest that these genes could be used as biomarkers for the diagnosing and monitoring the disease evolution.

**Figure3.**
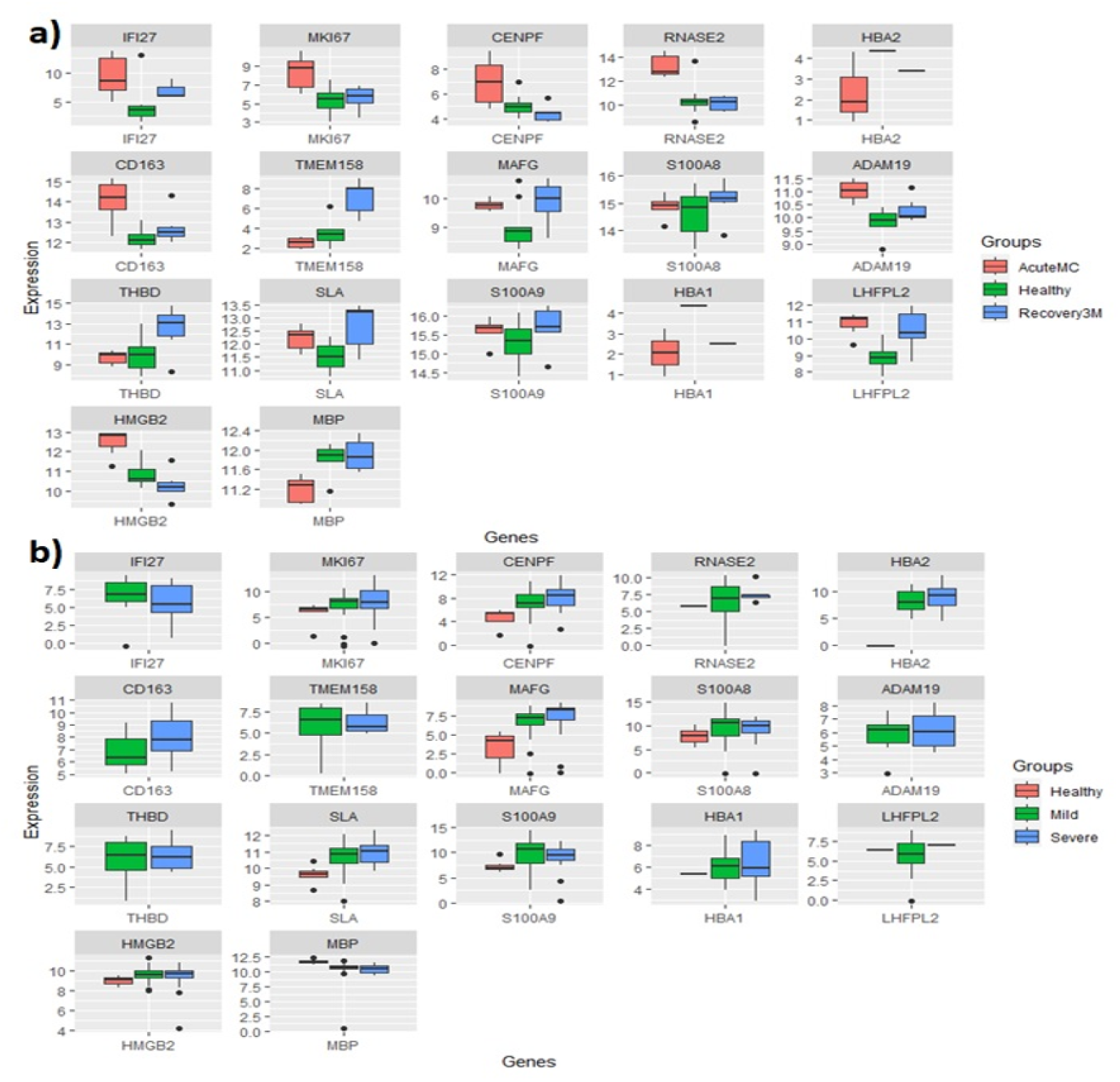
Genes Expression distribution: Expression distribution of *IFI27, MKI67, CENPF, RNASE2, HBA2, CD163, TMEM158, MAFG, S100A8, ADAM19, THBD, SLA, S100A9, HBA1, LHFPL2, HMGB2 and MBP genes* under different conditions of the two datasets GSE165461 dataset (NK) and GSE198256 dataset (MC). a) Boxplot of 17 genes expression levels across GSE198256 dataset (MC) three groups which are: 1-eleven controls “Healthy”, 2-seven COVID-19 patients “AcuteMC” and 3-Six samples of recovered people after 3 months from the infection “Recovery3M” b) Boxplot of 17 genes expression levels across GSE165461 dataset (NK) three groups which are:1-eight controls “Healthy”, 2-thirty nine COVID-19 patients with mild symptoms “Mild” and 3-twenty five COVID-19 patients with severe symptoms “Severe”.

The comparison of the expression levels of these genes among the nine datasets that were used in this study, as well as a complementary analysis of these genes are provided in supplementary information, and it includes the output of GeneALaCart application from the GeneCards online platform(S2) [47].

### Applying Filters for the Identification of Promising Biomarkers for Accurate COVID-19 Severity Assessment

Diagnostic strategies that are currently used for COVID-19 are mainly serological testing, antigen testing and nucleic acid testing, and despite their advantages, there is a need to identify additional targets for more accurate and efficient diagnosis of COVID-19 to handle the therapy based on the severity at the early stage of infection. In this regard, in the current study, we aimed at identifying potent markers for the early detection of the severity of the disease, indeed we added an additional filter based on the differential expression on the GSE165461 dataset (NK) between the severe patients and healthy donors sub-dataset as well as between mild patients and healthy donors sub-datasets. The different plots and the Venn diagram results are presented in Figure 4. Indeed, the results obtained showed 7 common dysregulated genes (*IFI27, MKI67, CD163, MAFG, ADAM19, SLA* and *MBP*) and the distribution of gene expression within the two datasets is depicted in Figure 3. More information on these genes and their expressions is presented in Figure-4.

**Figure-4.**
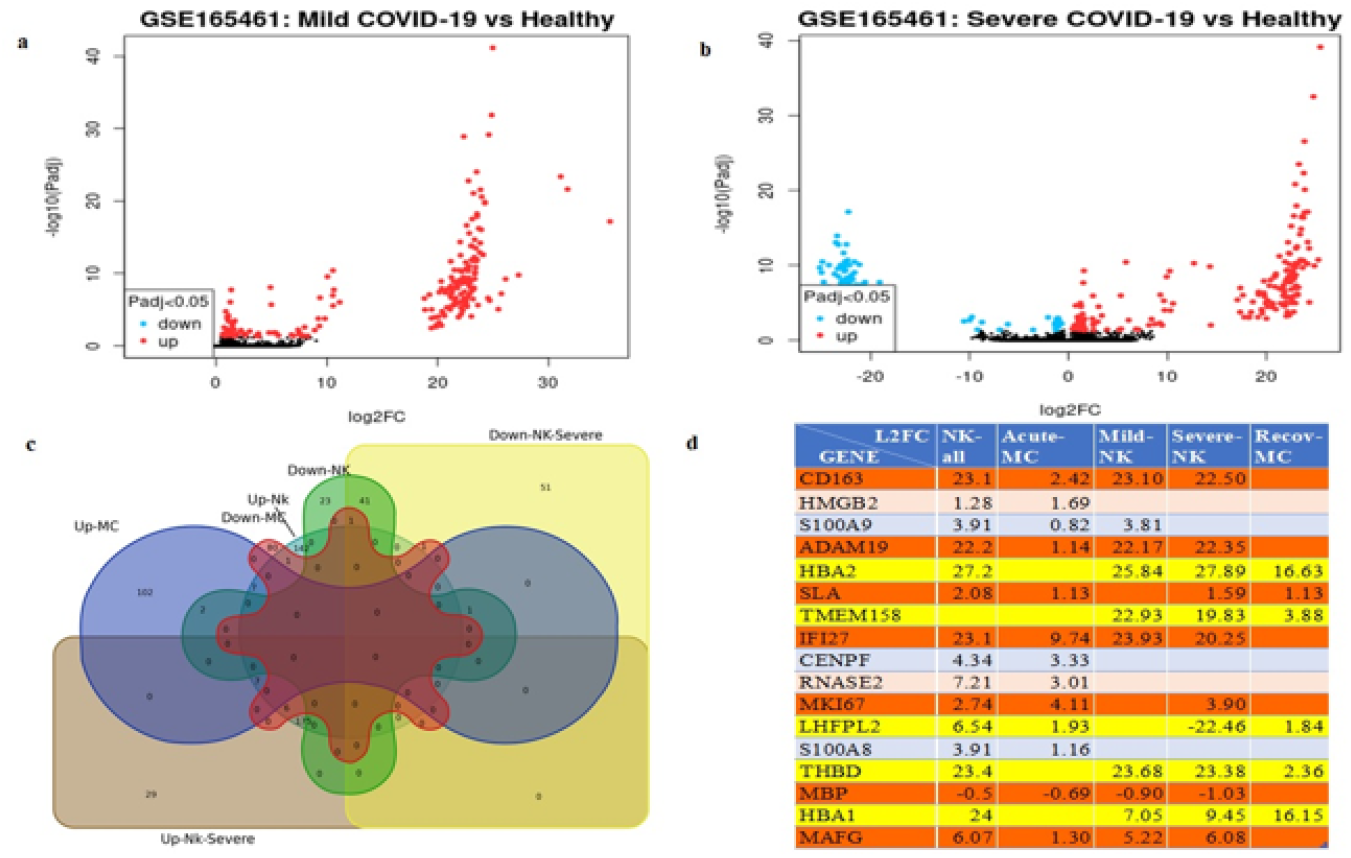
Transcriptomic signature of disease severity: Transcriptomic signature of disease severity diagnosis in COVID-19 patients: Identification of seven dysregulated genes and potential biomarkers, a&b) the volcano plot of GSE165461 dataset (NK) differential expression in the sub dataset of mild Covid-19 and the sub dataset of severe Covid-19, c,d) Identification of seven common dysregulated genes (*IFI27, MKI67, CD163, MAFG, ADAM19, SLA and MBP*) in the datasets GSE165461 dataset (NK) and GSE198256 dataset (MC) and their sub-datasets for disease severity diagnosis: A comparative Venn diagram and gene list analysis Venn, the red lines are for filtering per severity and the yellow ones correspond to the filtering per Long-Covid.

The common downregulated gene obtained was the *MBP* described earlier. The common up-regulated genes such as *IFI27, MKI67, CD163, MAFG, ADAM19, and SLA* play different roles in the regulation of cell proliferation, DNA replication, transcription, cell cycle progression, immune response, and inflammation. There isn’t much data on how the upregulation of these genes affects COVID-19 specifically, but some studies imply that inflammation- and immune-response genes like CD163 and MAFG may have increased expression as well. This suggests that severe COVID-19 may be linked to these genes. In fact, in a study analyzing gene expression profiles of COVID-19 patients, upregulation of *IFI27* was found to be associated with disease severity and poor prognosis[32]. Another study found upregulation of CD163, a marker of alternatively activated macrophages, in severe COVID-19 cases, suggesting a role in the development of hyperinflammation[34].

The results obtained from our analyses indicate that these seven genes could be considered as good predictors of the severity of the disease and may therefore be used for a precise and rapid diagnosis of COVID-19 severity and progression.

Further filtering can be performed based on the genes that remain overexpressed even after recovery. Among the 17 genes listed earlier, six genes continue to be overexpressed in individuals who have recovered from the infection after three months. These genes are *HBA2, TMEM158, THBD, SLA, LHFPL2 and HBA1. HBA2 and HBA1*, encoding hemoglobin subunits, demonstrate positive regulation, suggesting potential impacts on oxygen transport. *LHFPL2* plays a role in female and male fertility and is involved in the development of the distal reproductive tract[48], *TMEM158*, a transmembrane protein, may play a role in membrane-related functions, potentially influencing cellular processes. *THBD*, encoding thrombomodulin, exhibits heightened regulation, indicating a potential impact on blood coagulation. *SLA*, a Src-like adaptor, is also up-regulated, suggesting potential influences on cellular signaling pathways. These genes should undergo further evaluation to determine their potential application as biomarkers for long-term COVID-19.

### Applying Filters to Identify Promising Biomarkers for LONG COVID-19 Assessment

The condition where patients who have recovered from COVID-19 are still experiencing persistent symptoms after weeks or months is called Long COVID-19 [49], [50]. These symptoms may include shortness of breath, fatigue, chest pain, joint pain, sleeping difficulty and brain fog. What causes exactly long COVID-19 is not yet fully understood, but it is believed to be related to an overactive immune response or persistent viral infection in some cases [51].

We carried out a thorough literature review to confirm the potential of the six genes as predictors of long-term disease. We sought to examine the phenotypic capabilities of immune cells from people with long COVID-19 symptoms as part of this work, thus we obtained the GSE224615 dataset, which was just recently published. This dataset included blood samples from 27 LC and 16 non-LC participants from the San Francisco-based Long-term Impact of Infection with Novel Coronavirus (LIINC) cohort, and RNA-seq experiments were conducted on the blood specimens. The comparison of this dataset with our own yielded partial success: It revealed that 4 out of identified genes are commonly upregulated, with the addition of a new gene exhibiting upregulation. The five genes of interest are *HBA2, TMEM158, THBD, HBA1 and ANKRD33B (Ankyrin Repeat Domain 33B)*, that warrant further investigation. (S3)

The *HBA2* and *HBA1* genes have been identified as critical players in oxygen transport from the lungs to various peripheral tissues. Moreover, research has demonstrated that these genes function as antagonist peptides of the cannabinoid receptor CNR1. Hemopressin-binding to *HBA2 and HBA1* blocks the CNR1 receptor and subsequent signaling, which has implications for long-term diseases such as chronic pain, anxiety, and depression, where the CNR1 receptor has been implicated[52].

The *TMEM158* gene has been the subject of many studies exploring its potential role in various biological processes and diseases. Research has revealed that *TMEM158* can behave like a tumor suppressor gene and play a role in the mutator pathway of colorectal carcinogenesis. Moreover, TMEM158 has been identified to play a role in neuronal survival. Furthermore, studies have suggested that this gene can be used to diagnose anaplastic thyroid carcinoma and it has been also reported to promote pancreatic cancer aggressiveness by activating PI3K/AKT and TGFβ1 signaling pathways. On the contrary, *TMEM158* may also promote the proliferation and migration of glioma cells through STAT3 signaling in glioblastomas. Overall TMEM158 may have a complex role in different types of cancers and potentially serve as a therapeutic target in certain diseases[53].

Multiple articles explore the *THBD* gene, focusing on its role in regulating blood clotting and its connections to respiratory diseases. Topics include the impact of oxidized low-density lipoprotein on thrombomodulin transcription, thrombomodulin’s involvement in ARDS and SARS development, and the association between soluble thrombomodulin levels and mortality in ARDS patients. One article also reveals a BioID-derived interactome for SARS-CoV-2 proteins, highlighting protein-protein interactions in living cells [54].

*ANKRD33B* (Ankyrin Repeat Domain 33B) is a protein coding gene. There is limited information on the pathways and biological processes where this gene is involved although the diseases associated with *ANKRD33B* include Human Granulocytic Anaplasmosis.

Both *ANKRD33B* and *TMEM158* genes are also over expressed in GSE166253 dataset within retested positive (RTP) covid-19 patients.

A complementary analysis of these genes is provided in supplementary information and it includes the output of GeneALaCart application from the GeneCards online platform (Fishilevich et al., 2017).

Additionally, a long non-coding RNA (lnc-RNA) gene was commonly upregulated and may deserve further studies. In fact it’s about the gene *AC104564*.*3* which is very poorly described in literature; and recently, it has been found to be involved in a diagnosis and therapeutic responses models of immune-related lncRNA Pairs in clear cell renal cell carcinoma[55]

On the other hand and to broaden our analysis of GSE198256, we extended our study to include datasets from 6 months post-recovery, alongside the initial 3-month data. This expanded scope allowed us to capture a more comprehensive view of the long-term recovery process from COVID-19, enhancing our understanding of the disease’s enduring impacts. Our findings from this broader dataset revealed significant gene expression changes that persisted or emerged between these two recovery phases, providing deeper insights into the molecular recovery trajectory.

By tracking gene expression at 3- and 6-months post-recovery from COVID-19, we identified several genes with significant regulatory changes that provide insights into the complex recovery dynamics. TMEM158 and ANKRD33B, for instance, showed notable upregulation, highlighting their roles in cellular signaling and immune system recalibration. In contrast, MBP’s decreased expression could suggest lingering neurological impacts, common in long COVID. Additionally, genes like CD163 and THBD, which were upregulated, indicate ongoing inflammation control and vascular repair processes. These findings not only deepen our understanding of the post-COVID-19 physiological landscape but also suggest new therapeutic targets and biomarkers to enhance long-term recovery strategies. This holistic view of gene expression changes offers a robust framework for further research and clinical applications aimed at improving outcomes for COVID-19 survivors.

we utilized the ExpressAnalyst platform [56] to perform a comprehensive enrichment analysis on the curated list of genes identified above. These genes, along with their respective log2 fold changes, were analyzed against integrated databases including KEGG, Reactome, and Gene Ontology categories (Biological Processes, Molecular Functions, and Cellular Components). This analysis helped in mapping the genes to relevant biological pathways and functions, facilitating the identification of statistically significant enrichments that could provide deeper insights into the physiological and molecular mechanisms influenced by COVID-19.

Figure 5 showcases detailed network visualizations that bring to light several key biological domains influenced by COVID-19. The visualization highlights significant changes in cellular architecture and signaling, particularly noting alterations in membrane components that could impact cell signaling and structural integrity. The analysis also draws attention to the neurological impact of the virus, revealing enriched pathways involving neuronal structures like axons, which align with current findings on COVID-19’s neurological effects. Additionally, the study sheds light on the disease’s effect on blood coagulation processes, notably hemostasis and the coagulation cascade, aligning with the clinical signs of coagulopathy observed in patients. Furthermore, it highlights notable shifts in molecular functions such as ion and peptide binding, suggesting changes in cellular interactions that could influence metabolism and cellular response mechanisms. The analysis also details pathways associated with immune and inflammatory responses, including the AGE-RAGE signaling pathways and complement systems, providing vital insights into the inflammatory dynamics that are pivotal in understanding COVID-19’s extensive impact on human health.These insights not only augment our understanding of COVID-19’s broad systemic impact but also highlight potential molecular targets for therapeutic strategies. By delineating the complex interplay of biological processes altered in the disease, this analysis provides a foundational framework for further investigation into pathophysiological mechanisms and therapeutic interventions.

**Figure-5.**
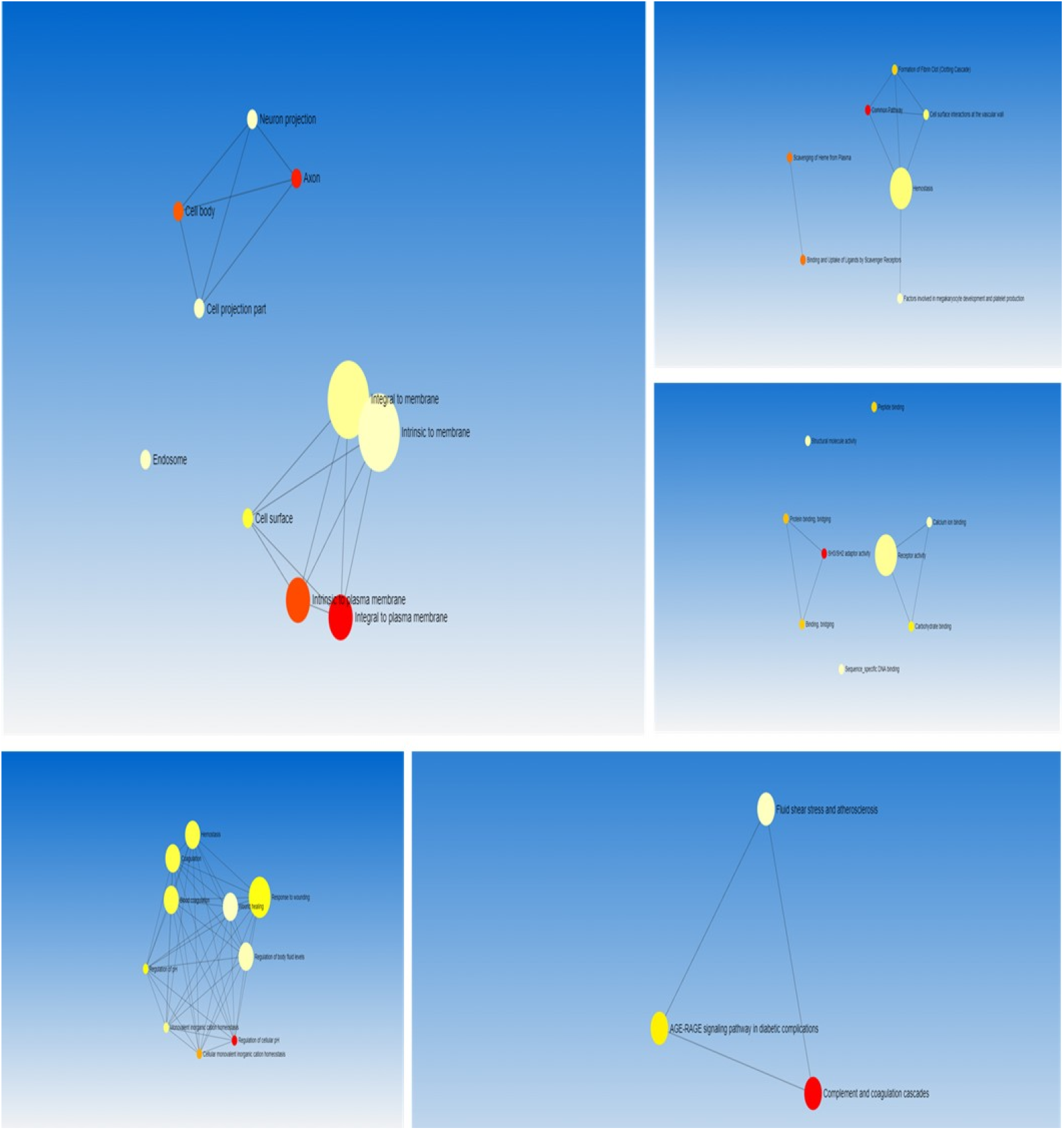
Comparative Network Analysis: Comparative Network Analysis of Dysregulated Genes Between 3-Month and 6-Month COVID-19 Recovery Phases

Moreover, the significant downregulation of IFI27 in the 6-month samples compared to healthy controls points to a diminished antiviral response, which might impact patients’ immunity long after the acute phase has resolved. This finding is crucial as it highlights potential vulnerabilities in recovering patients and underscores the importance of monitoring and possibly enhancing viral defense mechanisms during post-COVID care.

### Cross-Validation of Identified Biomarker Candidates

To explore the significance of the previously identified genes, we incorporated two additional datasets into our literature review. The first dataset, GSE152418, deals with RNAseq analysis of peripheral blood mononuclear cells (PBMCs) from 17 COVID-19 subjects and 17 healthy controls. Arunachalam PS, Wimmers F, Mok CKP, Perera RAPM et al. analyzed this dataset in their study published on September 4, 2020[57].

The second dataset, GSE215262, focuses on transcriptional patterns of PBMCs in SARS-CoV-2 infected patients with varying severity and outcomes. Qian X, Wu B, Chen X, Peng H et al. conducted the analysis in their study published in June, 2023[58].

Our results are promising, with many of the selected genes showing dysregulation in both datasets. Notably, *HBA1, HMGB2, HBA2, S100A9, TMEM158, IFI27, CENPF, RNASE2, MKI67, and S100A8* are consistently overexpressed in COVID-19 patients across both datasets. Additionally, *MBP* consistently exhibits downregulation among COVID-19 patients in GSE152418 datasets, suggesting potential significance in the context of COVID-19. (Figure-6)

**Figure-6.**
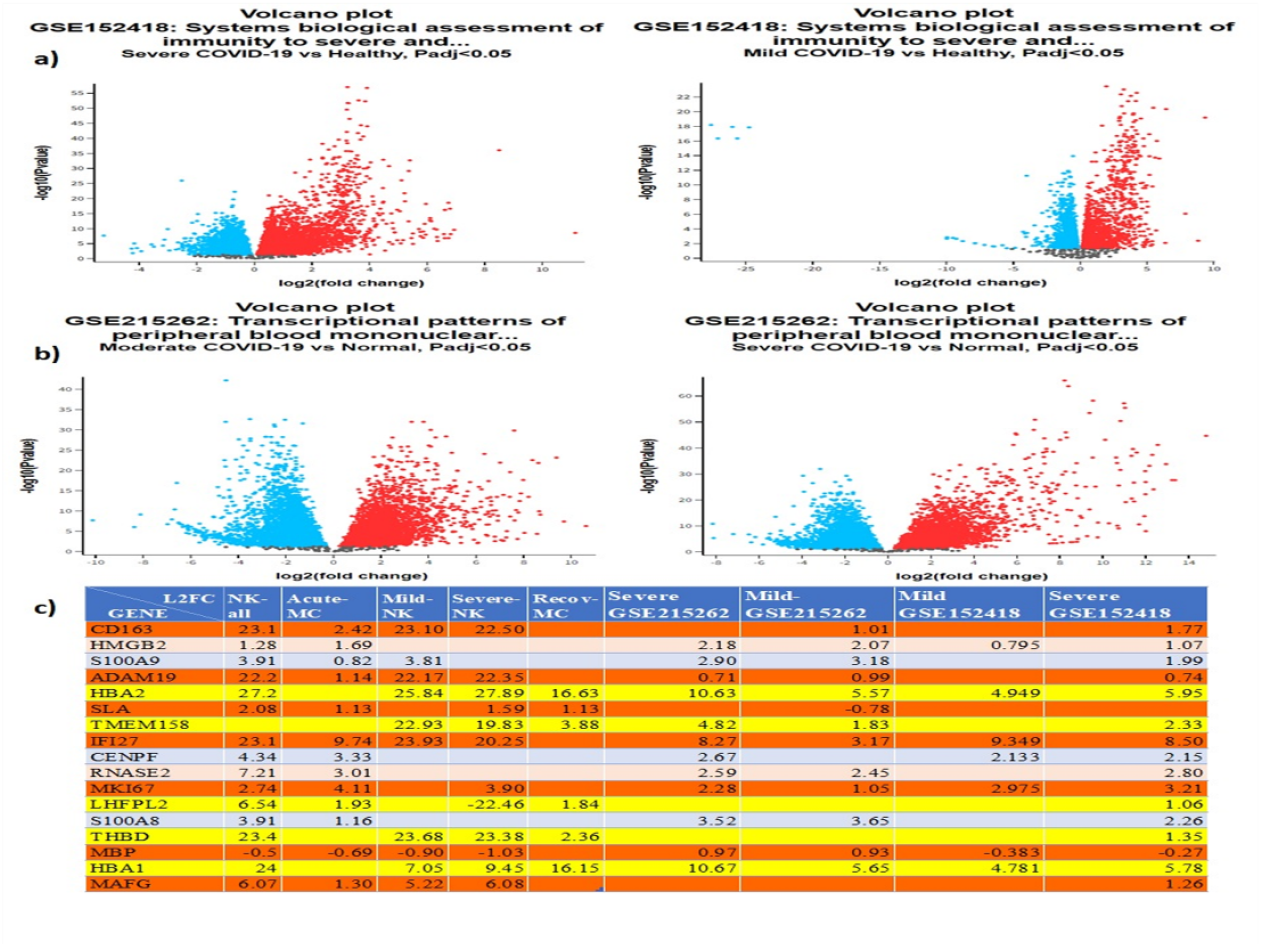
Transcriptomic profiling of COVID-19 for cross validation: a,b) Volcano plot of GSE152418 dataset and GSE152418 datasets as per their sub dataset of Moderate and Severe Covid-19, c)a list of the common dysregulated genes and their log2fold change expressions.

### Review on known drugs and regulators for the common dysregulated genes

Two gene analysis tools, GeneAlaCart[47] and GeneAnalytics[59], were used to gain valuable insights into potential drugs and compounds that could regulate the identified dysregulated genes. The results of the analysis are accessible at https://ga.genecards.org/shared-query/a0ec35e6-3169-4d36-9044-50ca8f00fb84.

To expedite the testing process, we filtered for approved bioactive drugs. Once more, we investigated the importance of the genes identified earlier as therapeutic targets by analyzing their expressions in two additional datasets, GSE182917 and GSE151973.

The GSE182917 dataset includes transcriptome sequencing data from lung tissue of 11 COVID-19 patients and 3 individuals with other lung pathologies, representing “Lung” data. This dataset compared the immune profiling expression patterns in COVID-19 decedents. The GSE151973 dataset includes biopsies collected through nasal endoscopic surgery from 4 adult patients, with samples harvested from both nasal respiratory epithelia and olfactory sensory epithelia. Bulk tissue RNA was extracted, libraries were generated, and the data is represented by “OE”.

The dysregulated genes identified in these datasets suggest that these genes may be involved in the entry of the virus into the lung tissue, nasal respiratory epithelia, and olfactory sensory epithelia tissues.

The common dysregulated genes with these two datasets include: *IFI27, MKI67, CENPF, RNASE2, CD163, LHFPL2, HMGB2*; *S100A8, ADAM19*, and *S100A9*..

The approved drugs and known regulators that may target these genes are filtered to include only bioactive compounds where at least three genes are involved. The list of identified compounds are presented in Figure-7. More information are available in the supplementary information in the UnifiedDrugs sheet within the GeneAlaCart file where more details are provided on their role and action as well as papers (PubMed IDs) which associate the gene with the drug.

**Figure-7.**
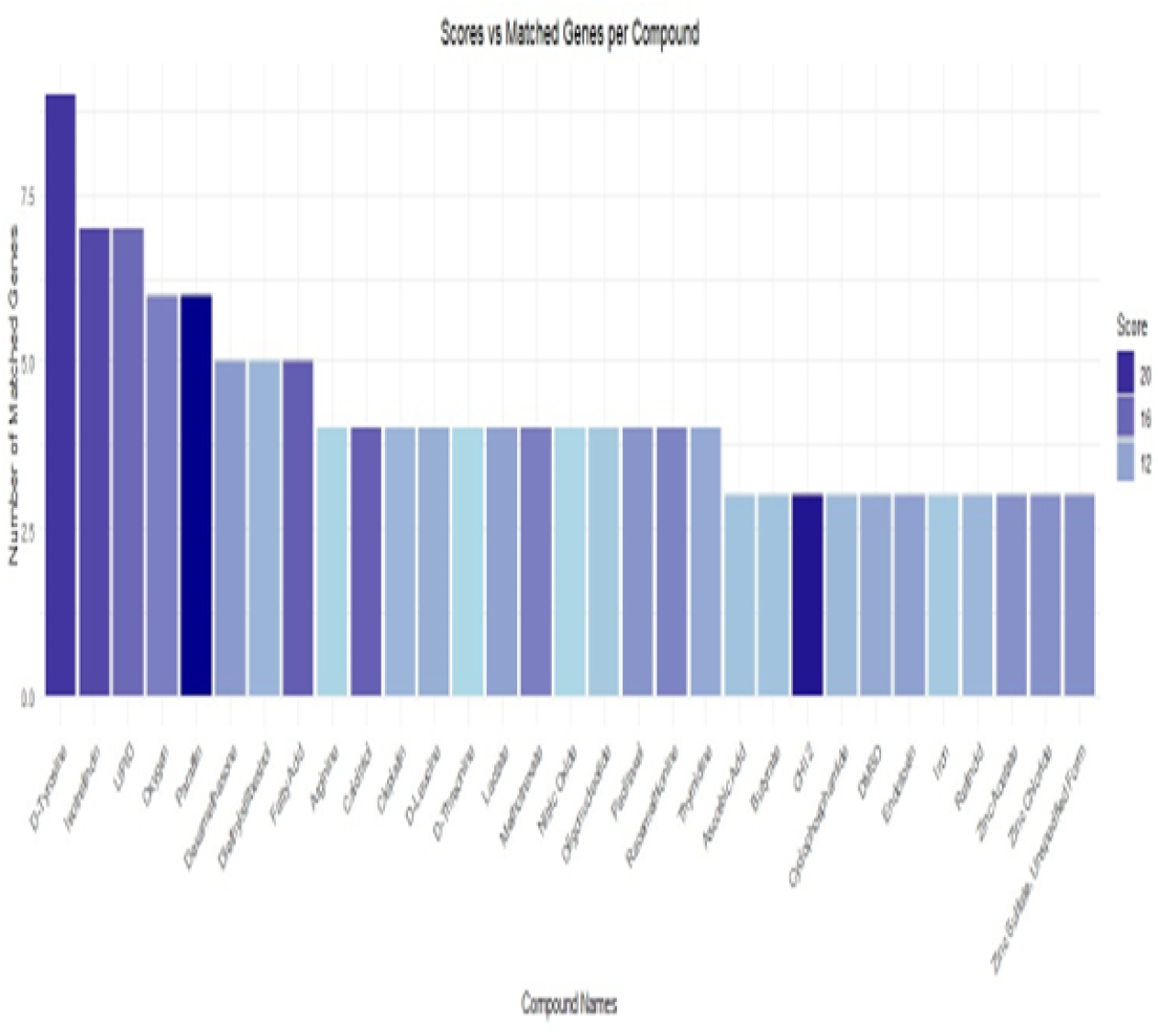
The list of potential therapeutic compounds: The graph of list of compounds with the number of genes that may be involved.

Further investigation into these medications is necessary to conduct a comprehensive analysis, considering the conditions they aim to address, potential side effects, recommended dosages, and the individual characteristics of the patients. Nevertheless, certain drugs on the list may exhibit fewer side effects or generally be well-tolerated, Table-1.

**Table-1:**
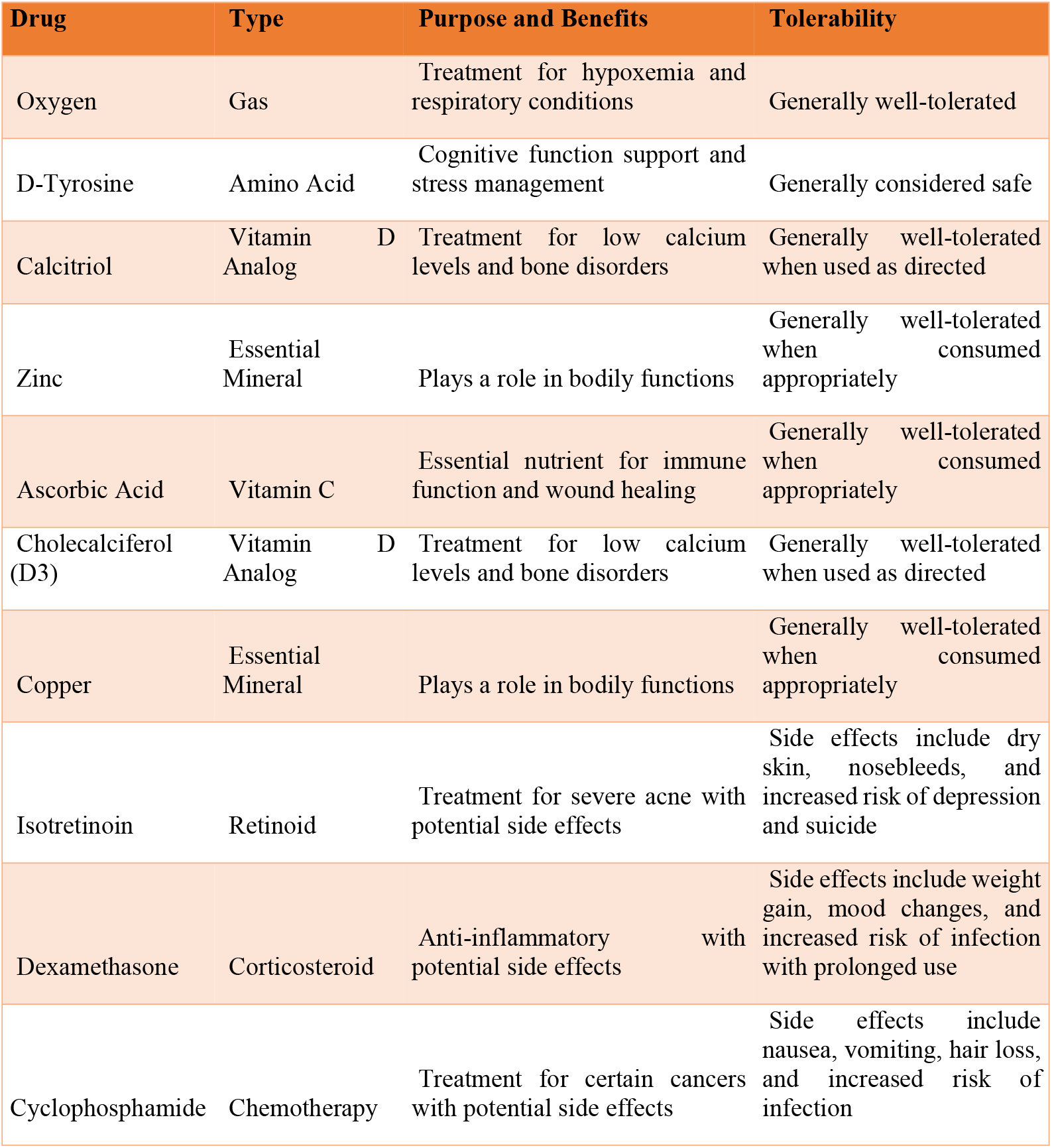
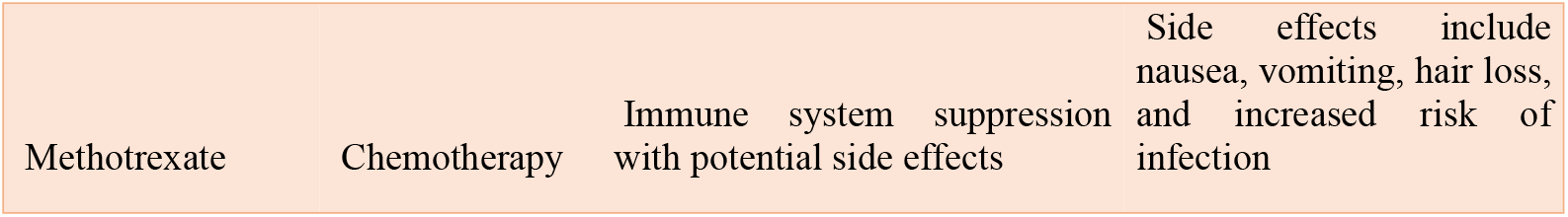
Selection of Possible Compounds and their general purpose and tolerability.

It is important to note that just because a drug may have a lower incidence of side effects or a better safety profile than another does not mean that it is always the best option for a certain condition or specific patient. The decision to use a particular drug should always be based on a careful evaluation of the potential benefits and risks, taking into consideration the patient’s medical history, current health status, and other relevant factors. However, in-depth investigation into this issue may assist patients and healthcare professionals in making knowledgeable judgments regarding available treatments.

## 4 Material and Methods

### Data Collection and Preprocessing

For our study, we selected publicly available gene expression datasets from the Gene Expression Omnibus (GEO) that focus on COVID-19 in human subjects. These datasets were chosen because they provide insights into the immune responses from four specific tissue types—peripheral blood mononuclear cells (PBMCs), lung, olfactory and sensory epithelium (OE), and lymph. These tissue types are critical in the immune system’s reaction to infections and are potentially valuable in identifying diagnostic and therapeutic targets. To maintain a focus on standard physiological responses, our selection excluded datasets from single-cell studies, cell lines, or those involving pharmacological treatments. In the preprocessing phase, we retrieved raw count data and gene annotations directly from GEO. We implemented a filtering step to ensure data quality, keeping only genes with a minimum of ten counts in the smallest sample group. This approach was designed to ensure that subsequent analyses were based on robust and representative gene expression data.

### Data Normalization and Differential Expression Analysis

In this phase of our study, we utilized the DESeq2 package [13], a widely respected tool within the bioinformatics community, for normalization and differential expression analysis. This tool helped us adjust for variations in library size and compositional biases that are typical in sequencing data. We conducted our analysis using two statistical tests: the likelihood ratio test (LRT), which provided a comprehensive ranking of genes, and the Wald test, which we used to explore specific contrasts between groups. To maintain rigorous statistical integrity, we applied the Benjamini-Hochberg procedure to control the false discovery rate [14], setting a stringent adjusted p-value cutoff at 0.05. Furthermore, to ensure that our findings were reproducible and transparent, we utilized GEO2R, an online tool available on the GEO platform. This tool allowed us to validate our results and provides a pathway for other researchers to replicate our analysis and verify our findings easily. Each dataset underwent this thorough analysis process, confirming the statistical robustness and reliability of the identified differentially expressed genes.

In anticipation of future studies, we are considering the adoption of alternative techniques that offer varied approaches to normalization and variance estimation. Notably, edgeR [15], which excels in managing small sample sizes and effectively handles overdispersion in count data, and limma-voom[16], originally designed for microarray data but now adeptly modified for RNA-seq analyses requiring robust linear modeling. Moreover, we aim to explore cutting-edge RNA-seq tools like kallisto | bustools, STARsolo, Salmon Alevin, GLM-PCA, and scVI, available through the Gene Expression Omnibus. These tools are particularly effective in processing complex, large-scale, and single-cell datasets, crucial for delving into the nuanced details of transcriptomic data. For example, kallisto | bustools and Salmon Alevin streamline transcript quantification, enabling rapid processing of extensive data ([17], [18], while GLM-PCA and scVI offer advanced methods for uncovering hidden data patterns and integrating diverse datasets [19], [20]. These approaches are invaluable for generating deeper insights into the disease mechanisms and cellular responses integral to conditions like COVID-19. Employing these innovative technologies not only advances our understanding but also assists in identifying new diagnostic markers and therapeutic targets, expanding the potential applications of GEO datasets in translational medicine.

### DEG files Review and verification

The preliminary findings from various datasets align consistently with existing literature and published papers, reinforcing our strategies for identifying potential targets that have not yet been experimentally tested. These datasets cover diverse aspects of COVID-19 pathology, ranging from immune responses in specific cell types to the exploration of distinct clinical conditions. Notably, our analysis sheds light on dysregulated genes associated with conditions such as severe COVID-19 and long COVID-19, providing valuable insights into the underlying immunological features. The following table highlights key studies associated with each dataset, offering a comprehensive overview of the dysregulated genes and their implications (Table-2).

**Table-2:**
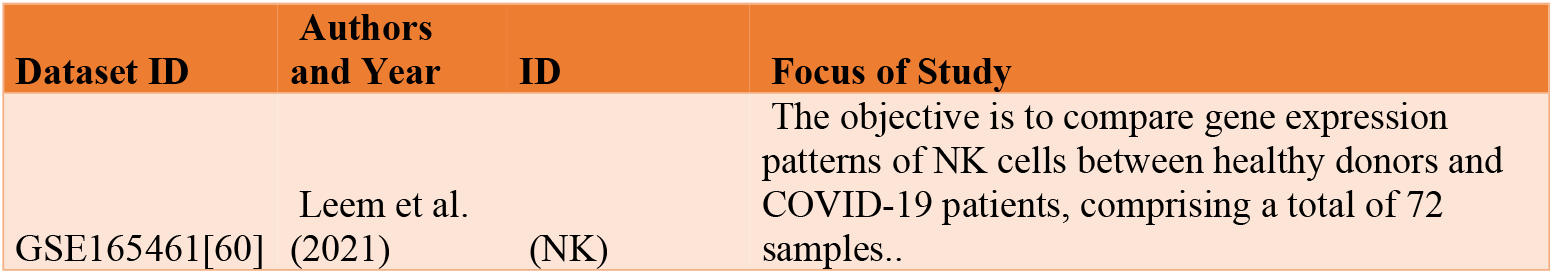
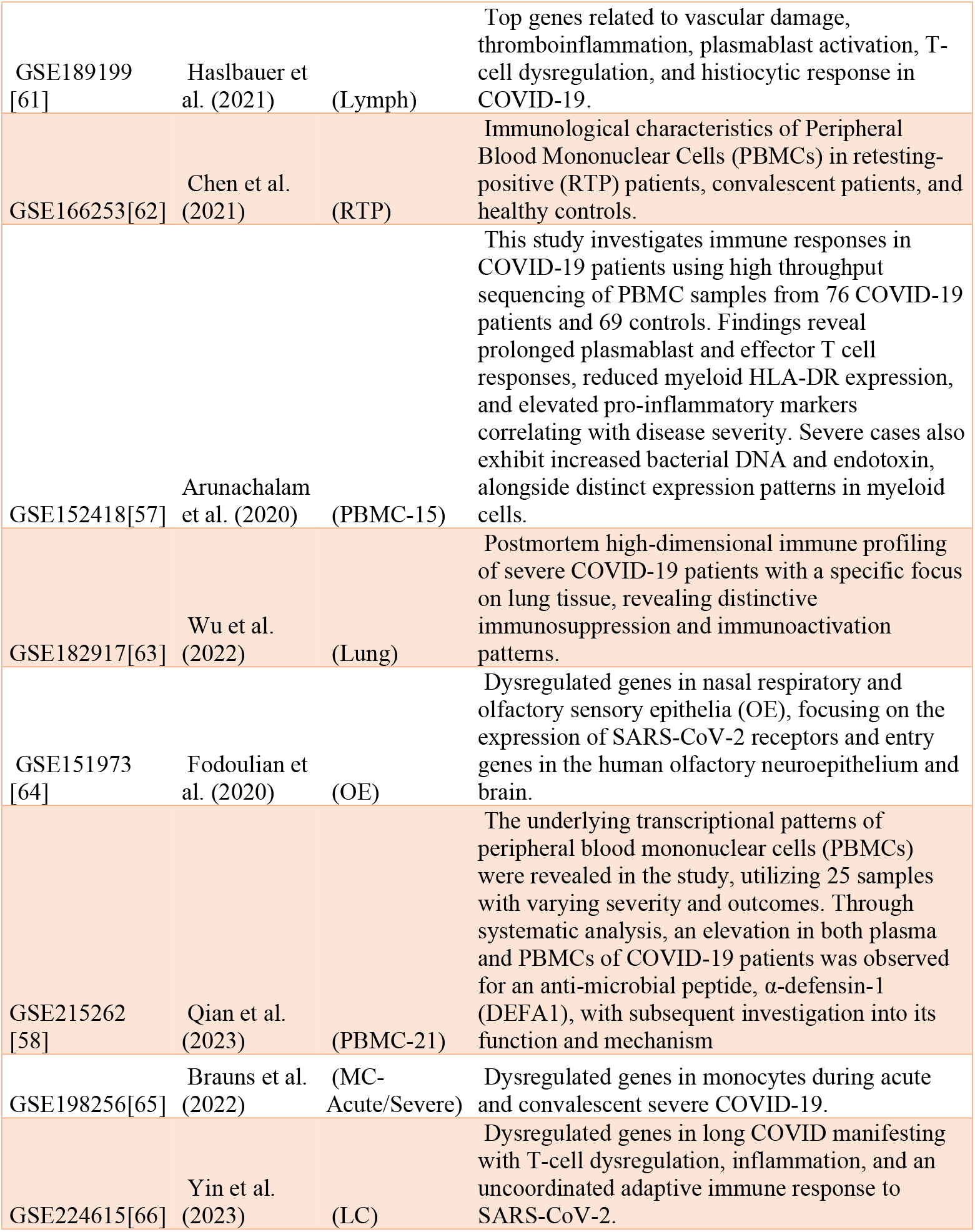
Published studies on the datasets used.

### Integration of Differentially Expressed Genes

We synthesized the differentially expressed genes from each dataset into a unified list, focusing on those consistently dysregulated across varied datasets. This methodical integration was crucial for pinpointing robust gene candidates that hold potential both as diagnostic biomarkers and as targets for therapeutic intervention. Specifically, we categorized these genes into two key groups: Biomarker Genes: These are pivotal genes identified as likely to enhance the accuracy and speed of COVID-19 diagnostics and Target Genes: These genes stand out as promising candidates for the development of innovative therapeutic strategies.

### Functional Enrichment and Pathway Analysis

To delve deeper into the biological implications of the identified dysregulated genes, we utilized a suite of enrichment analysis tools including GeneCards [21] Gene Ontology (GO), and the Kyoto Encyclopedia of Genes and Genomes (KEGG). These tools helped us to map out the biological pathways impacted by these genes, offering a clearer understanding of how they might influence both the progression and the potential treatment options for COVID-19. This analysis is crucial for pinpointing specific biological processes that could be targeted to mitigate the disease or enhance therapeutic strategies Our approach to presenting the results included a range of visualization techniques that not only illustrate but also simplify the understanding of complex data. Volcano plots were particularly useful in highlighting genes with significant expression changes, offering a visual contrast between magnitude of effect and statistical confidence. We have detailed our analytical workflow in Figure 1.

### Validation Through Literature Review and Cross-Checking in Additional Datasets

To validate our findings, we conducted a comprehensive literature review, comparing our identified differentially expressed genes with previously published data to confirm their relevance and significance in the context of COVID-19. This step was crucial in establishing the credibility of our biomarker and therapeutic target identifications, ensuring that they are consistent with known biological mechanisms and previous research findings. Furthermore, to strengthen our conclusions and mitigate the risk of dataset-specific biases, we cross-checked our results against additional datasets from the Gene Expression Omnibus (GEO). This included analyzing similar tissue types and clinical conditions to see if the dysregulated genes we identified were consistently expressed in other independent studies. This dual approach of literature validation and empirical cross-dataset verification provides a robust framework for confirming the potential of these genes as reliable markers for diagnosing and treating COVID-19, thereby enhancing the translational value of our research.

## 5 Conclusion

This study embarked on a detailed exploration of the Gene Expression Omnibus (GEO) datasets to uncover potential biomarkers for COVID-19, emphasizing the pivotal roles of NK cells and CD14+ monocytes during the 3- and 6-month recovery phases and the severity stage of the disease. Utilizing the analytical power of DESeq2, we identified significant genes such as *TMEM158, ANKRD33B, LHFPL2, SLA, CD163, MAFG, MBP*, and *THBD* that were differentially expressed across these time points, highlighting their potential as key players in both the diagnosis and longitudinal monitoring of the disease.

Our findings reveal a landscape where these biomarkers not only shed light on the immediate response to infection but also on the prolonged biological processes that govern recovery and possibly the persistence of symptoms known as Long COVID. For instance, the ongoing upregulation of *TMEM158 and ANKRD33B* may reflect a sustained immune activation, potentially impacting long-term health outcomes. Conversely, the consistent downregulation of MBP could suggest lingering neurological impacts, aligning with the cognitive challenges observed in Long COVID patients.

Moreover, our comparative analysis further identified a distinct expression pattern in tissues directly involved in the viral entry, like lung and nasal epithelia, underscoring genes that could serve as targets for early therapeutic intervention. This not only enriches our understanding of the disease’s pathology but also opens up new avenues for targeted treatments that could mitigate the initial viral assault and subsequent complications.

To further our understanding, we recommend incorporating multi-omics approaches in future research. This would not only enhance the depth and resolution of our insights but also provide a more comprehensive view of how these biomarkers function within the complex network of human biological processes. Additionally, leveraging biobank samples for empirical validation of our findings could help substantiate and expand upon our results. Such rigorous validation is essential for refining diagnostic techniques and tailoring treatment strategies to better meet individual patient needs. Ultimately, these steps will enhance the precision of patient management and care, contributing to more effective health outcomes in response to this global health challenge.

Moreover, the approach of integrating and comparing diverse, heterogeneous datasets, as employed in this study, offers a valuable framework not only for understanding COVID-19 but also for addressing other diseases. This methodology enables the identification of shared genetic mechanisms and the discovery of novel therapeutic targets across various conditions. Applying this approach to other diseases in the future could significantly broaden our understanding of complex biological processes and improve our capacity to address public health challenges with greater accuracy.

In summary, our research highlights the crucial role of robust genomic data and advanced data analysis in uncovering the complex dynamics of COVID-19 recovery. By identifying key biomarkers and clarifying their roles throughout the disease’s progression, this work contributes to the broader efforts to combat COVID-19 and establishes a foundation for managing future public health crises with greater precision.

## Author Contributions

Conceptualization and methodology, SL, RBM, NB, REL and RB; Data collection and analysis, SL and, RBM, writing—original draft preparation, SL and RBM; supervision, RBM, NB, REL and RB; All authors participate in writing, reading and correcting the manuscript and agreed to its contents.

## Declaration of generative AI and AI-assisted technologies in the writing process

During the preparation of this work AI-assisted technologies were used to improve the readability and language of the manuscript. After using this tool, the authors reviewed and edited the content as needed and take full responsibility for the content of the published article.

## 1 Supporting information

1-S-Supplementary files, a word document including:

- Datasets description.
- The table of the aggregated DEG files in an embeded excel Sheets
- Log2Fold Change of the identified genes expression across the different datasets
- Publication mini-review on the gene TMEM158
- Publication mini-review on the implication of the THBD gene in respiratory diseases
- GeneAlaCarte files in an embeded excel Sheets

2-GEO2R Scripts in a Zip file

